# Stressful exposure from the 2004 Indian Ocean Tsunami drives long-term changes in thyroid hormone physiology

**DOI:** 10.64898/2026.09.17.752410

**Authors:** Ralph Lawton, Duncan Thomas, Elizabeth Frankenberg, Cecep Sumantri, Teresa Seeman, Eileen Crimmins, Daniel Hochbaum

## Abstract

Stressful events are associated with long-term adverse health impacts, but the causal mechanisms linking these exposures to subsequent disease remain poorly understood. One plausible mechanism is persistent changes to hormonal signaling that regulate metabolic homeostasis. Here, we examine the long-term impacts of stress associated with exposure to the 2004 Indian Ocean tsunami on levels of free triiodothyronine (FT3), a circulating measure of biologically active thyroid hormone, using data from the Study of Tsunami Aftermath and Recovery. We measure biomarkers 20 years after the tsunami in a population-representative sample age 35y+ in 2024 who, at the time of the tsunami, were living along the coast of Aceh, Indonesia, the most affected part of the Indian Ocean Basin. Because respondents were first interviewed before the tsunami and subsequently tracked regardless of migration, the sample avoids a major source of selection bias common in long-term disaster studies. We identify the causal effect of tsunami exposure using variation in community-level tsunami mortality, comparing communities within the same sub-district. We find that greater tsunami exposure reduced levels of FT3 20 years later. Exposure also altered relationships between FT3, body composition, and cortisol and increased cardiometabolic risk. These findings identify persistent alteration of thyroid hormone physiology as a potential pathway linking severe stressful exposures to long-term cardiometabolic disease risk.

**Significance Statement:** Exposure to severe stressful events is associated with worse cardiometabolic health, but the biological mechanisms underlying these long-term effects remain poorly understood. Using the 2004 Indian Ocean tsunami as a natural experiment, we show that greater tsunami exposure causally reduces circulating free T3, the biologically active thyroid hormone, two decades later. Tsunami exposure also alters the normal relationships between free T3 and other metabolic measures and increases cardiometabolic risk, suggesting that persistent reductions of free T3 in response to stress may contribute to adverse cardiometabolic outcomes. Together, these results identify persistent changes in thyroid hormone physiology as a previously underexplored response to severe stressful exposure and a potential pathway linking such events to long-term metabolic disease.

## Introduction

Exposures to events causing extreme stress are common and harmful to health (Chauntry et al. 2026; Cohen, Janicki-Deverts, and Miller 2007; Hikichi et al. 2021). Understanding the mechanisms through which stress affects physical health in the long term is critically important but methodologically complex, especially in human populations (Adam 2023; Hansen et al. 2025; Miller, Chen, and Parker 2011). Strikingly, circulating hormone levels can change with exposure to a major stressor. Given their control of metabolic processes throughout the body, changes in hormonal set points could drive the downstream consequences for metabolic health commonly observed in populations exposed to chronic stress. For instance, elevated cortisol after a stressor becomes maladaptive if it persists over the longer term, resulting in adverse health outcomes (Russell and Lightman 2019) and, in some cases, ‘cortisol burnout’ (Lawton et al. 2023; McEwen 2007; Takahashi et al. 2021). Here, we examine the role of stress on long term levels of thyroid hormone, which is a key regulator of metabolism, suggesting a pathway through which stress may affect cardio-metabolic health risk (Abdalla and Bianco 2014). Thyroid hormone actions are widespread, for example, regulating LDL-receptor expression and cholesterol clearance in the liver, adipocyte differentiation and lipid mobilization in fat cells, and substrate metabolism in skeletal muscle (Brent 2012; Mullur, Liu, and Brent 2014).

System-wide, this has direct implications for blood sugar, cholesterol, body composition, and markers for cardio-metabolic disease (Han et al. 2015; Lawton, Sabatini, and Hochbaum 2024; Nam et al. 2010). Further, thyroid hormone has potent effects on behavior through direct effects on brain plasticity and circuitry (Hochbaum et al. 2024). Though a large literature links stress to cardiometabolic health, often implicating cortisol responses and the hypothalamic-pituitary-adrenal-axis (HPA-axis), the hypothalamic-pituitary-thyroid axis (HPT-axis) is likely critical yet has received much less attention (Cohen et al. 2012, 2007; Hikichi et al. 2021; Kivimäki, Bartolomucci, and Kawachi 2023; Lawton et al. 2023; Seeman et al. 2018; Takahashi et al. 2020).

Despite the importance of thyroid hormone for metabolic health and behavior, and strong evidence of disruption caused by stress in the short term, the long-term response of the HPT-axis to stress has not been established (Friedman et al. 2005; Jung et al. 2019; Machado et al. 2015; McEwen 2004; McIver and Gorman 1997; Nadolnik 2011; Wang and Mason 1999). The core circulating HPT-axis cascade consists of Thyroid Stimulating Hormone (TSH), secreted by the pituitary gland, thyroxine (T4) secreted by the thyroid, and triiodothyronine (T3) – the primary active hormone of the chain, produced through conversion of the prohormone T4. In this paper, we focus on T3 because it is the primary ligand for thyroid hormone receptors and binds them with approximately 10-fold greater affinity than T4, thereby mediating most canonical thyroid hormone-dependent transcriptional effects (Abdalla and Bianco 2014). In the short-term, stressors like hospitalizations, acute physical stress, or inflammatory changes are associated with well-documented decreases in circulating thyroid hormones, often called non-thyroidal illness syndrome (Chatzitomaris et al. 2017; Petrowski and Kahaly 2025). The HPA-axis may in part mediate these short-term responses, as cortisol alters thyroid hormone production, but the long-term relationship is less clear (Helmreich et al. 2005; McEwen 2004; Petrowski and Kahaly 2025). Prior work on long-term relationships between stress and thyroid hormones in humans is limited, typically conducted with small and selected samples, with inconsistent results regarding associations between earlier life stressors and thyroid hormone levels – some studies find increases and others find decreases in T3 and other thyroid hormones (Friedman et al. 2005; Jung et al. 2019; Mason et al. 1994; Toloza et al. 2020).^1^ Because isolating a causal effect of stress exposure on thyroid hormone levels in human populations is a substantial challenge, most scientific evidence is based on animal models (Petrowski and Kahaly 2025). To address these challenges we analyze the long-term impacts on thyroid hormone levels of a specific, well-identified, stressful exposure. Specifically, we treat exposure to the 2004 Indian Ocean tsunami in Aceh, Indonesia, as a natural experiment, and use purpose-collected population-representative biomarker data to compare FT3 levels 20 years later among individuals with differing exposure. This study provides rigorous evidence to establish the impacts of stress on FT3 levels over the long term in human populations. We extend these analyses to examine relationships between FT3 and other biomarkers, including body fat and cortisol, to characterize the potential directionality of the relationships between thyroid function and other health markers after stress. Our analyses identify lowered FT3 levels, and by extension, dysregulation of the HPT-axis, as a likely pathway through which long-term metabolic outcomes are affected by stress.

Our approach measures the physical health impacts of exposure to the stresses of a large-scale natural disaster, the 2004 Indian Ocean tsunami. Using data from the Study of the Tsunami Aftermath and Recovery (STAR) collected for this project, we measure individuals’ exposures to stress from the tsunami by constructing the tsunami-related mortality rate in each of the 96 communities in which individuals resided at the time of the tsunami and draw comparisons between adults (35y+ at time of measurement, 15y+ in 2004) from nearby communities to identify a dose-response impact of stress on FT3 levels measured 20y post-tsunami. Since the tsunami was completely unanticipated and the community-level tsunami mortality rate is quasi-random conditional on geographic characteristics, we are able to isolate the long-term impacts of exposure. The research design addresses three major historical limitations on studies of the long-term consequences of stress. First our measure of stressful exposure is objective and well-measured. This is rare; most studies rely on respondent recall of events that occurred many years in the past. Second, we establish that community-level tsunami mortality is plausibly exogenous to other drivers of later life health. Third, twenty years post-tsunami we measure biological parameters for a population-representative sample that was interviewed pre-tsunami and followed since early 2004 with high retention rates (>93%). This is important because the study sample is representative of all tsunami survivors and not selected on factors such as displacement and migration, which is likely related to the stress of the exposure. Many post-disaster studies rely on selective samples, which is a major limitation.

We find that long-term FT3 levels decrease as community mortality rate at the time of the tsunami increases and that the impacts are greatest at older ages. Cardio-metabolic risks also rise with increasing tsunami exposure. Our evidence suggests that changes in FT3 levels are an upstream driver of cardio-metabolic risk, and we are able to characterize tsunami-related disruption of canonical links between FT3, cortisol, and body fat. These results extend our prior work, which finds long-term impacts of tsunami exposure on the HPA-axis and on cognition (Lawton et al. 2023, 2025; Sheridan et al. 2024), and are, to our knowledge, the first report of HPT-axis output (FT3 levels) as a potential driver of long-term metabolic outcomes associated with stress.

Our results also contribute to the literature on long-term consequences of disaster exposure on prime-age adults, which is more limited than studies of those in utero or in older age groups at exposure (Barker 1994; Kawachi et al. 2020; Strauss and Thomas 1995). Our focus on the long-term impacts of stress on thyroid hormone yields insights into the mechanisms underlying the relationship between stress and cardio-metabolic health. Further, we find that the impacts of the 2004 tsunami emerge at older ages in particular, emphasizing the importance of long-term follow up to study the health impacts of events earlier in life.

## Results

### Sample population and comparisons with the US

Thyroid biomarkers were collected as part of a sub-study conducted on all tsunami survivors who were living (at the time of the tsunami) in randomly selected communities that were included in the STAR baseline. The Extended Assessments of Biomarkers and Cognition sub-study selected one in four baseline communities, oversampling communities that were hardest hit by the tsunami. Demographic characteristics of our analytic sample of 3,270 respondents aged 35 and older when the biomarkers were measured are displayed in Table 1. The average respondent is 50.9 years old, has 9.1 years of education and there are slightly more females than males.

**Table 1.** Summary statistics and exogeneity tests.

|  | [1] | [2] | [3] |
| --- | --- | --- | --- |
| Variable | Mean | Standard Deviation | p-value for regression on tsunami mortality |
| Age at survey | 50.9 | 11.6 | 0.508 |
| Male (%) | 49.2 | 50.0 | 0.729 |
| Height (cm) | 156.6 | 8.2 | 0.342 |
| Years of education, baseline | 9.1 | 3.6 | 0.664 |
| Baseline ln(HH per-capita expenditure) | 12.64 | 0.51 | 0.206 |
| Current smoker (%) | 31.5 | 46.5 | 0.578 |

Notes: STAR analytical sample: 3,270 individuals. Columns 1 and 2 report unadjusted descriptive statistics for the sample. Column 3 reports the relationship between characteristics and the treatment variable of interest: community-level mortality rates at the time of the tsunami (quartic root), estimated with sub-district fixed-effects, controls for elevation and distance to the coast, and with standard errors clustered at the community level.
<sup>3</sup> The monthly average high temperature in Aceh fluctuates between 86 and 90F during the year; in the US, it fluctuates between 67 and 85F between May and October. In NHANES, average FT3 levels are 0.1 standard deviations higher in the other months; free T4 levels are 0.03 standard deviations higher (not statistically significant) and TSH levels are the same, conditional on age, sex, and race (Lawton et al. 2024).
<sup>4</sup> For binary classifications, we use a damage zone classification that stratifies communities into those heavily damaged versus not heavily damaged, based on information sources that include satellite imagery, community informant reports collected after the tsunami, and the observations of interviewers working in the field after the tsunami. (Frankenberg et al. 2008)
<sup>5</sup> In the figures, the NHANES sample is restricted to those measured in May-Oct and in the age figure, the NHANES sample is restricted to non-obese (BMI<30) individuals to mitigate body composition differences across samples. These adjustments substantially affect the concordance between the NHANES and STAR demographic patterns. The NHANES FT3-age slopes are roughly 30% steeper, and FT3-BMI slopes for males are 50% steeper, without these adjustments.

To place the distributions of thyroid hormones in STAR in context, we compare them with those in the NHANES, a population-representative sample from the US which we age-and-sex-standardized to match the STAR sample (Appendix figure 1). Our primary analyte of interest is free T3 (FT3), which has been shown to better stratify and identify features of the thyroid axis in the general population, effects that are much smaller or absent in the upstream hormones of the thyroid signaling cascade TSH and FT4, and are attenuated somewhat for total T3 measures that combine FT3 and protein-bound T3 (Lawton et al. 2024). For budget reasons, we assayed other HPT-axis hormones, FT4 and TSH, for a smaller, randomly-selected sub-sample of the FT3 sample. (Abdalla and Bianco 2014; Lawton et al. 2024). Since FT3 levels are lower in higher ambient temperatures (Kuzmenko et al. 2021), the comparison is restricted to NHANES respondents surveyed in the summer months (May-October)^2^, when ambient temperatures in the US are closer to those in Aceh.^3^ FT3 levels in Indonesia are lower than those in the US, though the distributions overlap substantially and the standard deviations are very similar. The TSH distributions are nearly identical in STAR and NHANES and FT4 levels are substantially lower in STAR. Similar patterns of lower FT3, higher FT4, and similar TSH compared with the US have been observed in other countries, including India and Thailand, but population-representative data on thyroid hormone levels outside the U.S. is limited (Marwaha et al. 2013; Sriphrapradang et al. 2014). FT3 differences may reflect differences in body composition across contexts, as well as large differences in typical ambient temperature (Kuzmenko et al. 2021; Lawton et al. 2024).

**Figure 1.**
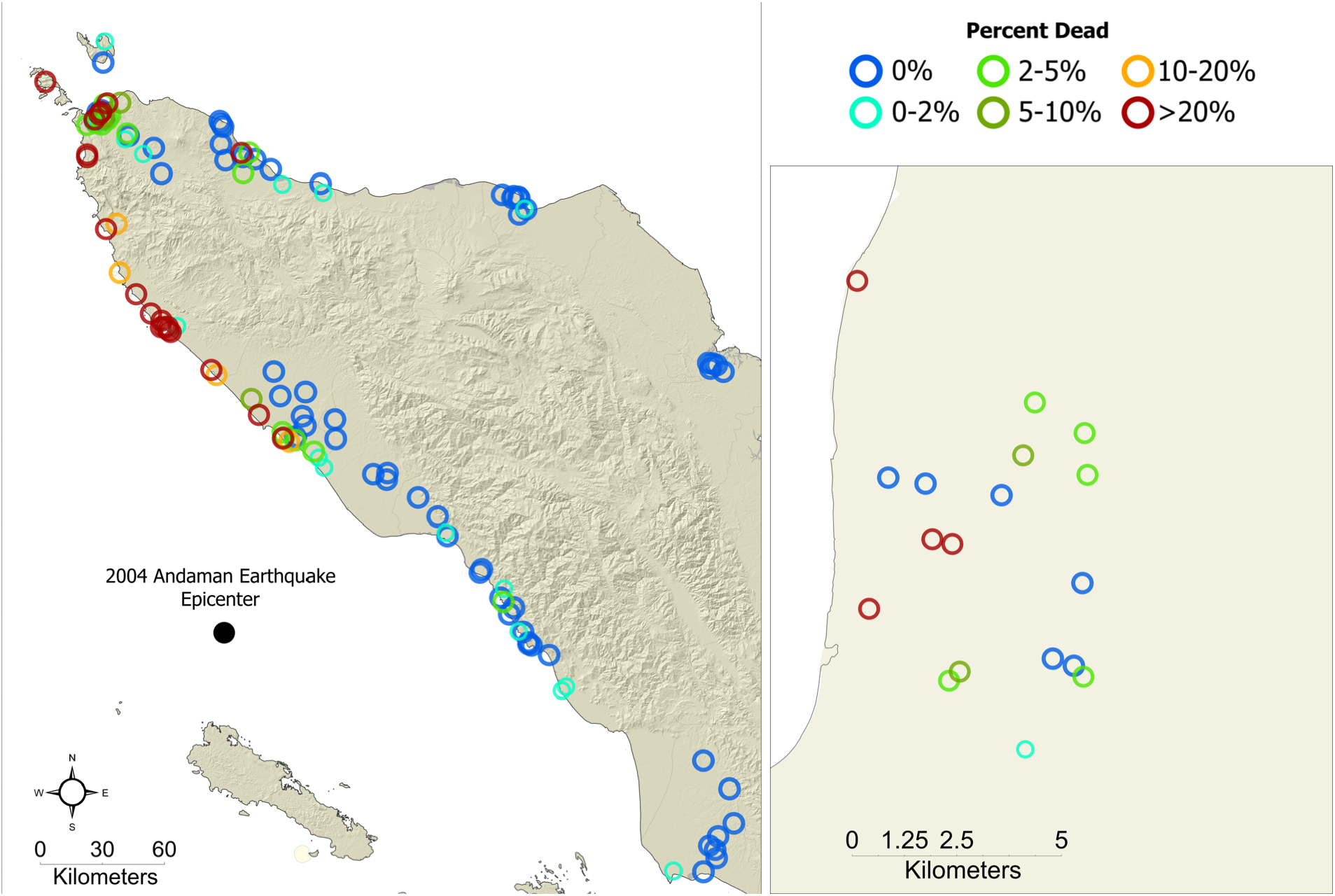
Baseline communities of residence in the STAR Enhanced Assessment of Biomarkers and Cognition sample. Colors denote the tsunami-related mortality rate in the community. Right panel magnifies a small area to illustrate distances between communities with different mortality rates.

Given the lack of population-level FT3 data outside of the US, we describe the demographic patterns of FT3 in our population. To limit contamination from tsunami exposure, we restrict the STAR sample for these relationships to those who were living, at the time of the tsunami, in communities that were not heavily damaged.^4^ Appendix figures 2 and 3 show the nonparametric BMI and age relationships with FT3. FT3 levels are expected to decline with age, and to have a positive association with BMI (De Pergola et al. 2007; Lawton et al. 2024). These patterns are observed for the STAR respondents. Males have higher levels than females, FT3 levels decline substantially with age, and are associated positively with BMI, mirroring relationships in the US. However, while the BMI gradients, and age gradients for females, are very similar in the two populations, the age gradients among males are not as steep in STAR as those in the NHANES.^5^ Taken together, we find qualitatively--- and, to a large extent, quantitatively--- similar demographic relationships with FT3 in the US and in the STAR sample.

**Figure 2.**
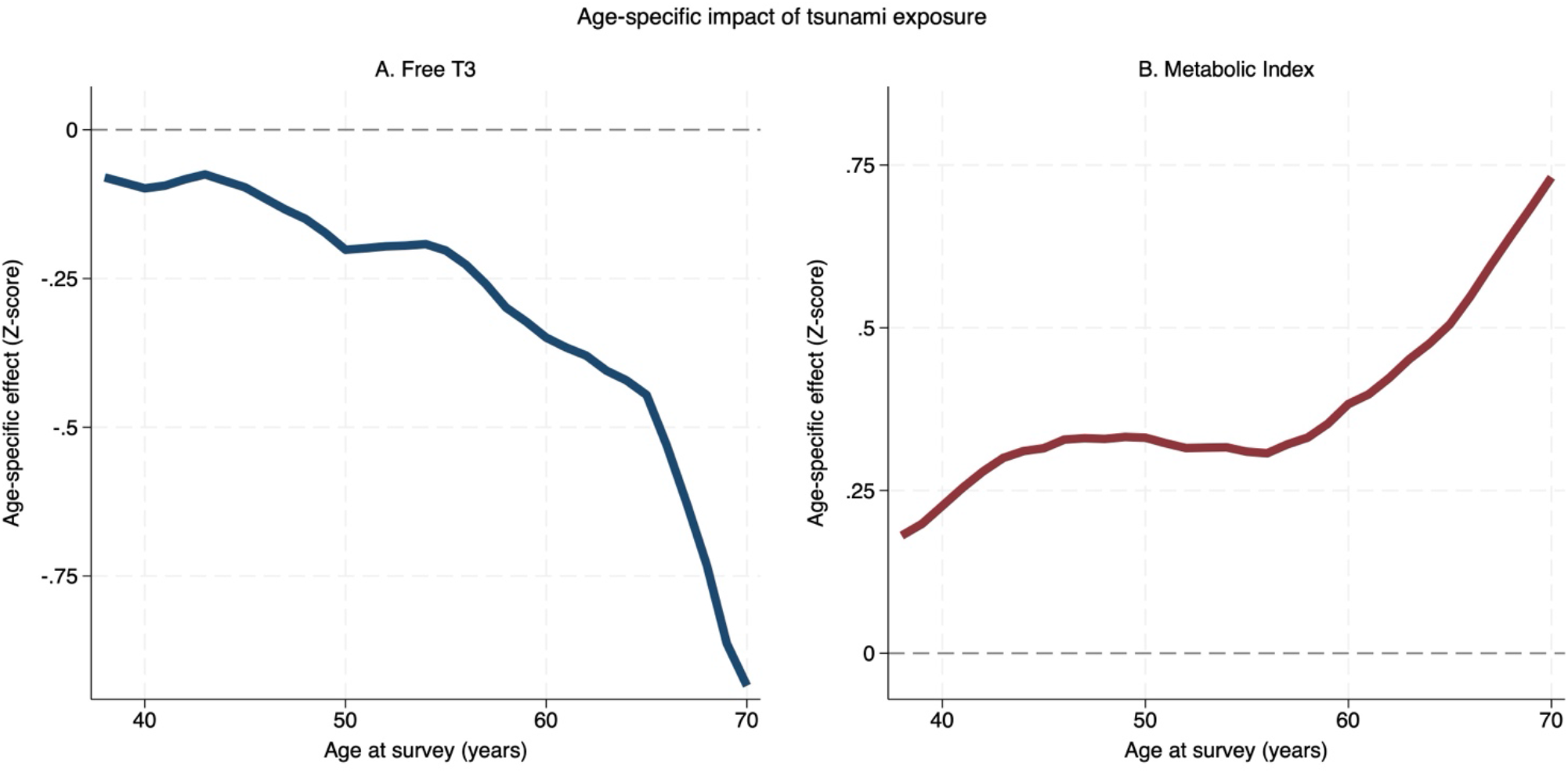
Age-specific estimates of the impacts of the tsunami on FT3 levels and the metabolic index. Estimates of the relationship between the quartic root of tsunami-related mortality in the baseline community are plotted for each age. Estimates are constructed by estimating the impact of tsunami exposure using observations surrounding a focal age – specifically, a bandwidth of 10 years is used alongside triangular weights, after both the exposure and the outcome were residualized on the covariates used to estimate the primary model in table 2.

**Figure 3.**
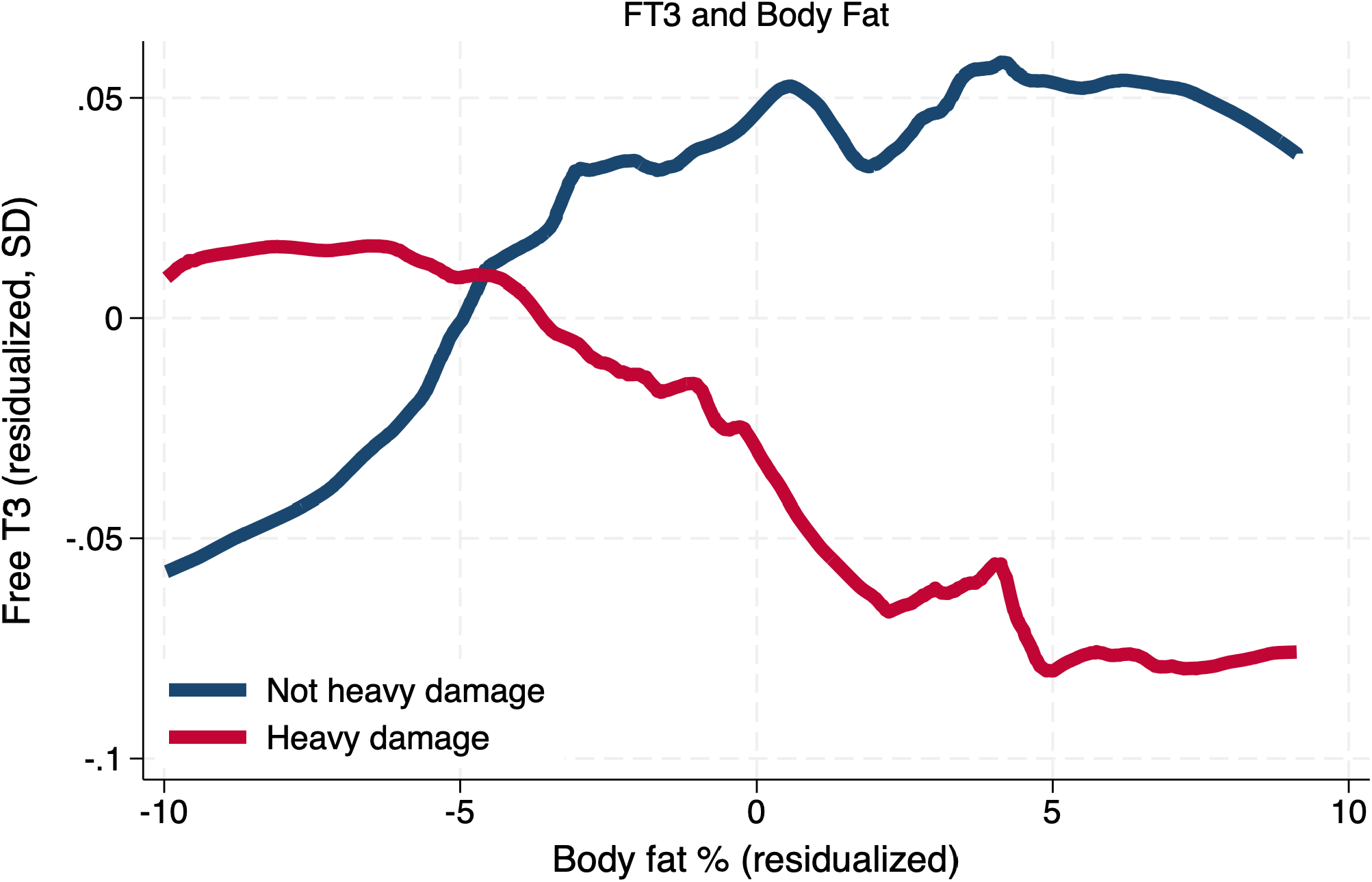
Non-parametric estimates of the relationship between body fat % and FT3 among individuals with varying tsunami exposure. We compare individuals who were living in communities with heavy tsunami exposure at baseline with those who were living in areas not heavily exposed. Stratified LOWESS estimates are presented, after residualizing the body fat percentage and FT3 levels on age and sex.

### Exposure to the 2004 Indian Ocean Tsunami

Our primary measure of tsunami exposure is the rate of mortality attributable to the tsunami in the community of residence at the time of the tsunami.^6^ The STAR study sites in which biomarkers were collected, color-coded by the tsunami mortality rate, are displayed in Figure 1. Local variation in exposure to tsunami mortality is substantial. One-third of the communities in this study had no tsunami mortality. In the other communities, on average 17% of the population died in the tsunami. Mortality is right skewed: in the median community with some mortality, 5% died and in the hardest hit community, over 75% died (Appendix figure 4). Even within a 5km area, mortality rates at the time of the tsunami varied dramatically (Figure 1, right panel). This local variation strengthens our identification strategy because we draw comparisons between people who were living, at the time of the tsunami, near to each other. By comparing people who were living in the same sub-district, we mitigate contamination in the estimates because of differences across geographies that includes, for example, socio-economic status, quality and accessibility of services. Since the distribution of mortality rates is right-skewed and includes zeroes, we use the quartic root transformation in the models (which approximates a logarithmic transformation for positive numbers).

**Table 2.**
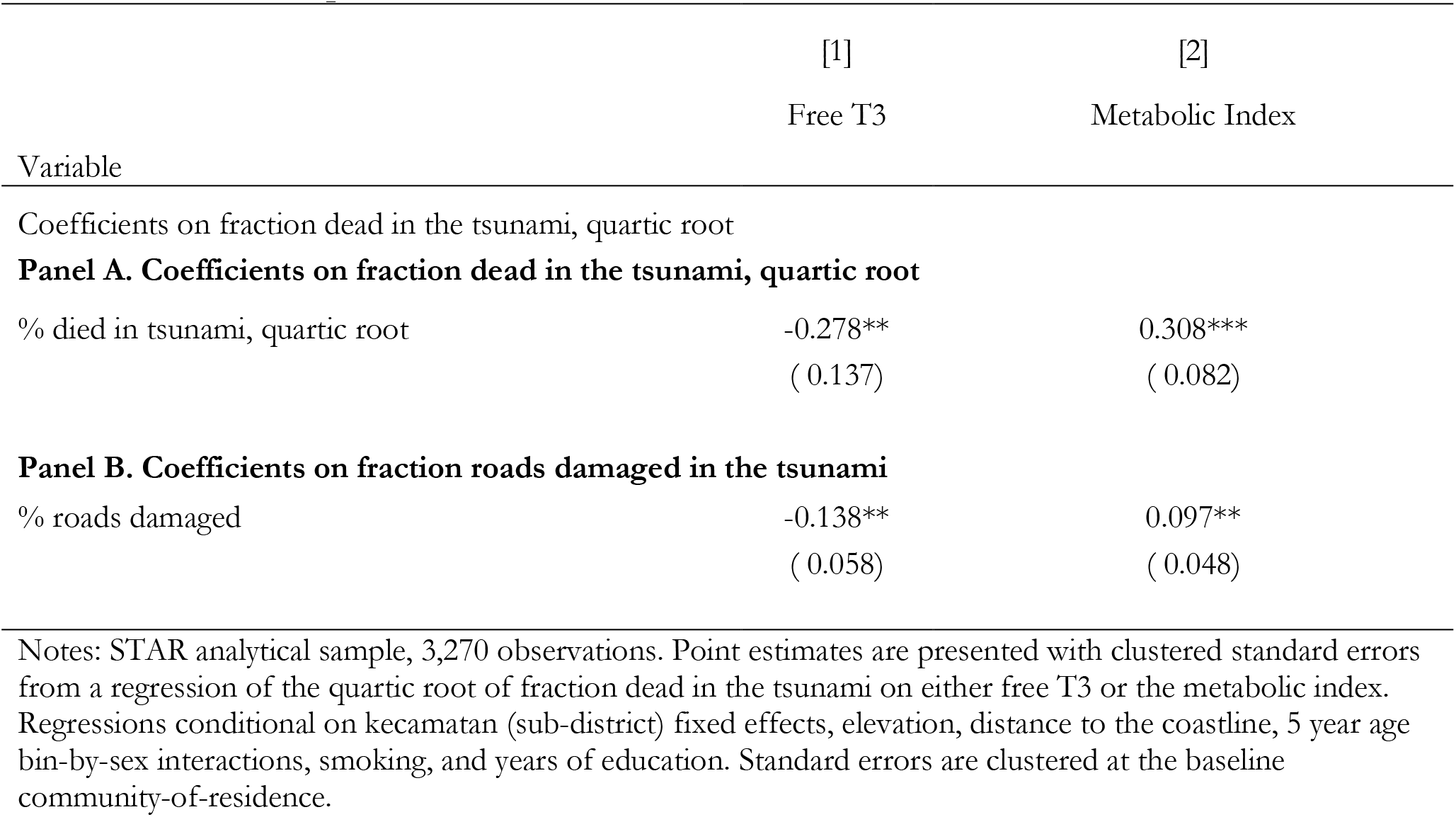
Tsunami exposure, free T3, and Metabolic outcomes.

Local variation in the tsunami’s lethality reflects differences across locations in the force and height of the waves, which in turn reflect features of the undersea rift zone that shifted in the earthquake and spawned the tsunami, in combination with the topography of the seafloor and coastline. Given the complexity of those relationships, in combination with the sudden and unanticipated nature of the tsunami, we treat variation in tsunami-related mortality rates within sub-districts as exogenous to future health. In support of this assumption, we show that tsunami exposure is unrelated to any of our baseline characteristics of interest (Table 1). Contemporaneous health behaviors, such as smoking, are also unrelated to exposure. Height, which in this context is an important indicator for early life health and current economic status, is unrelated to exposure (Deaton 2007; LaFave and Thomas 2017).

### Impacts of exposure to the tsunami on HPT-axis function 20 years later

In order to isolate the long-term impacts of the tsunami, we compare individuals from the same sub-district who were living in communities in which mortality rates from the tsunami differed, controlling for elevation and distance (as in column 3 of Table 1). Table 2 presents the *β* coefficients on FT3 expressed as a z-score (Panel A, column 1).

FT3 declines significantly as tsunami mortality increases (p=0.045). The magnitude of the decline is substantial: the implied impact of the difference in exposure between those in the least versus most affected communities is of similar magnitude to the difference between males and females, or the change in FT3 with over 20 years of aging.^7^ These findings provide causal evidence that exposure to a severe stressful event can produce long-term alterations in thyroid hormone physiology detectable two decades later. This is important because decreases in HPT-axis function have been connected to worse general health status and increased mortality risk, and the findings contribute to a small and mixed prior literature on the impacts of stressful events on thyroid hormone levels (Friedman et al. 2005; Jung et al. 2019; Mason et al. 1994; Toloza et al. 2020; Wang and Mason 1999). Further, this study provides this evidence in a population-representative sample with measurements of FT3, the principal bioactive thyroid hormone within the HPT-axis.

### Alternative measure of tsunami exposure

We complement our primary measure of tsunami exposure, tsunami-related mortality, with another continuous measure, the fraction of roads in the community that were destroyed by the tsunami. While both measures are continuous, and provide estimates of dose-response impacts, mortality is based on reported deaths in the study population and road damage is estimated by the community leader. As shown in panel B of Table 2, we draw the same conclusion that greater exposure reduces FT3. We draw qualitatively similar conclusions using our binary classification of damage zones but estimates are less informative than our continuous mortality measure as there is limited variation in the binary classification within sub-districts. These findings emphasize the value-added of the continuous measures of exposure.^8^

### Impacts of exposure to tsunami mortality on cardio-metabolic health 20 years later

Having demonstrated that stress affects FT3 levels, we next asked if there were similar impacts to cardio-metabolic health. We developed an inverse-covariance weighted index of cardio-metabolic disease measures: HDL and non-HDL cholesterols, HbA1c, systolic and diastolic blood pressure. We use this integrated index to address multiple-testing concerns (Anderson 2008). There is a significant increase in cardio-metabolic disease risk among those with more tsunami exposure, 20 years post-tsunami (p<0.001), and we find similar results when using fraction of roads damaged as an alternate measure of exposure (Column 2 of table 2). Cardio-metabolic risk is substantially worsened over the long term by tsunami exposure.

### Impacts of the tsunami exposure on long-term outcomes by age

Prior work examining the tsunami’s impacts on health, including post-tsunami mortality and cognitive functioning, have found that adverse impacts of tsunami exposure vary with age (Frankenberg, Sumantri, and Thomas 2020; Lawton et al. 2025; Sheridan et al. 2024). We find similar age-specific impacts in this study, both for the FT3 and the metabolic index outcomes. Figure 2 shows a non-parametric estimate of the age-specific effect of tsunami exposure on these outcomes 20 years later. The impact on FT3 is generally negative, but the estimated impacts grow as age increases. The impact of age on the metabolic index is positive, but here too the impacts at younger ages are much smaller than at older ages. For health characteristics that decline substantially with age, such as metabolic characteristics examined in this study, an intriguing hypothesis is that tsunami exposure shifts individuals on to a different trajectory of aging, suggesting that adverse health outcomes will continue to emerge as the cohorts age. Additional longitudinal data will be key for testing this hypothesis.

### Understanding the direction of impacts on HPT-axis function and metabolic health

Body composition, metabolic health markers, and FT3 typically exist in a regulatory equilibrium, but it is possible that exposure to stress produces changes in FT3 that drive downstream changes in metabolic conditions. We approached testing this hypothesis in several ways.

FT3 and body mass have a circular relationship and are usually in equilibrium: Positive energy balance and rising adiposity typically upregulate FT3 to increase metabolic rate, but exogenous increases in FT3 lower adiposity. This intuition underpins the simplest evidence that FT3 changes may be upstream of metabolic disease changes — the sign of the tsunami’s impacts on FT3 compared with sign on the metabolic index. Decreases in FT3 without decreases in the metabolic index suggest that FT3 is not responding to other changes in metabolic health or caloric intake (Biondi 2010; De Pergola et al. 2007; Rotondi, Magri, and Chiovato 2011). The pattern — lower FT3 coexisting with increased cardiometabolic disease risk — is inconsistent with FT3 responding to worsening metabolic health in equilibrium, and instead suggests that FT3 suppression may be contributing upstream to the other health changes we observe.

A second test finds suggestive evidence for a distortion of typical equilibria between thyroid hormone levels and body composition. We plot the nonparametric FT3-fat percentage relationships among individuals living in communities with and without heavy damage exposure, as classified by our binary measure of community-level tsunami exposure. Note that this figure uses a dichotomous measure of exposure rather than a continuous measure that isolates a dose-response. Figure 3 shows a clear difference in the relationship between those minimally exposed and those heavily exposed to the tsunami. For control individuals the relationship between FT3 and body fat % is positive, as expected. In contrast, for individuals in high exposure groups the relationship is negative, consistent with changes in FT3 that are exogenous rather than driven by a process of equilibration. In sum, these analyses suggest that FT3 levels could be a driver of broader cardio-metabolic health changes that have been documented in the aftermath of stressful events.

### Free T3, cortisol, and exposure to the tsunami

Acute stressors and activation of the HPA-axis have been canonically associated with depressed HPT-axis function (Helmreich et al. 2005; Petrowski and Kahaly 2025). The impacts of longer-term chronic stressors and PTSD, are less clear (Petrowski and Kahaly 2025). In a previous wave of STAR, we collected hair cortisol measures in a subsample of respondents, finding that tsunami exposure led to long-term reductions in cortisol, consistent with “burn-out” of the HPA-axis from prolonged activation (Adam 2023; Lawton et al. 2023; Miller, Chen, and Zhou 2007). Of those 630 individuals, 525 were surveyed in this wave, enabling a unique opportunity to examine the relationships between outputs of the HPA and HPT axes, and exposure to the 2004 Indian Ocean Tsunami. In Table 3, we examine the relationship between cortisol levels and FT3, using linear regression with interaction terms for tsunami exposure, such that the ln(hair cortisol) coefficients can be interpreted as the slope among those without direct tsunami exposure.

**Table 3.** Cortisol, thyroid hormone levels, and tsunami exposure.

| Variable | [1] | [2] |
| --- | --- | --- |
|  | Free T3 |  |
| ln(hair cortisol) | -0.192**<br>(0.08) | -0.172**<br>(0.07) |
| ln(hair cortisol) *<br>quartic root % dead in<br>tsunami | 0.211**<br>(0.09) |  |
| ln(hair cortisol) *<br>heavy damage<br>exposure |  | 0.117**<br>(0.05) |

Notes: STAR cortisol subsamples, for 525 respondents with hair cortisol measures at the 14 year wave who also have thyroid samples. Natural log of cortisol measures used (see: Lawton et al., 2023). Point estimates are presented with robust standard errors that show the linear relationship between the log of hair cortisol levels and free T3 in standard deviations, along with interactions by baseline community tsunami mortality rates or baseline community damage zone classification.

Among individuals without direct tsunami exposure, we find that we can replicate the canonical finding: cortisol levels and thyroid hormone levels are negatively related. However, we find that tsunami exposure severs this link: individuals with more tsunami exposure have much more positive relationships than those without. This is consistent with a long-term dysregulation of the HPA-axis by the tsunami, and with chronic changes in thyroid physiology that may not be contemporaneously mediated by the HPA-axis. The disrupted relationship between cortisol and FT3 levels suggest that stressors can lead to long-term remapping of hormonal homeostasis, and that the changes observed in FT3 are not simply downstream of contemporaneous HPA-axis disruption.

## Discussion

Twenty years after the 2004 Indian Ocean Tsunami those most exposed to its negative impacts exhibit different patterns of thyroid hormone physiology and metabolic health than those who were largely spared. Our results provide novel causal evidence of long-term impacts of stressful exposure on FT3 levels, complement other studies of the impacts of stress on cardio-metabolic health, and suggest potential mechanisms for how stressful events in adulthood impact later life health (Cohen et al. 2007; Kawachi et al. 2020; Kivimäki et al. 2023; Lawton et al. 2023, 2025; Seeman et al. 2018; Shiba et al. 2021).

Prior work has documented the long-term impacts of stress on the HPA-axis and related impacts on immunologic changes (Danese and McEwen 2012; McEwen 2012; Seeman et al. 1997). While HPT-axis function plays a central role in the regulation of metabolism, the impacts of stress on thyroid hormone levels has received much less attention, and prior literature studying FT3 is mixed. Studies of PTSD have found both hyper- and hypo-thyroid associations (Friedman et al. 2005; Jung et al. 2019; Machado et al. 2015; Toloza et al. 2020). A core conceptual challenge with studies that compare individuals with PTSD to those without PTSD, however, is the difficulty separating whether or not T3 levels are contributing to the presentation of or are resulting from PTSD (Toloza et al. 2020). Our emphasis on a specific, exogenous event rather than on a PTSD diagnosis directly rules out that T3 levels are endogenous to our exposure of interest, enabling causal interpretation of the finding that the tsunami led to long-term FT3 declines.

While the precise mechanisms through which HPT-axis changes may drive cardiometabolic disease require further study, the observed directions of effects and the altered FT3–body composition relationship in high-exposure individuals together suggest that subclinical HPT-axis changes likely drive broader deterioration of cardiometabolic health. This finding is of substantial importance. Prior work has suggested that stressful events and the responses to stress may shape cardio-metabolic disease (Hikichi et al. 2021; Seeman et al. 1997, 2018; Takahashi et al. 2020).

However, direct understanding of the potential mechanisms underlying these changes has been less clear. Long-term disruptions in the HPA-axis are one key pathway that may mediate these impacts (Lawton et al. 2023). The HPT-axis has received much less attention, despite its central role in regulation of metabolism and links to cardio-metabolic conditions. In our study, we find that canonical links between cortisol and FT3 levels are themselves shaped by tsunami exposure, suggesting that while stress may impact both the HPA and HPT axes and shape their long-term trajectories, contemporaneous effects of cortisol do not explain the long-term changes in FT3. This finding raises important mechanistic questions about the links between the HPT and HPA axes in the response to stress – in particular, the importance of chronic stressors, the downstream consequences of HPA-axis “burnout” that may accompany extended stressful events, and the time courses over which endocrine responses to stress may evolve. The interaction between these hormonal systems is a rich area for future work, and these findings suggest that thyroid hormone physiology warrants more attention in the long-term study of stress and health, both in human populations and in mechanistic studies of other organisms.

Our findings also underscore the importance of long-term follow-up. The results echo other long-term adverse health findings we have observed in aging-related markers including memory and HPA-axis dysregulation that persists over the long term (Lawton et al. 2023, 2025; Sheridan et al. 2024). The age-related patterns we observe suggest that these impacts may continue to emerge, and ongoing follow-up of these individuals will be essential for fully understanding the unfolding impacts of stress earlier in life.

There are several potential limitations to this study’s interpretation. First, without longitudinal markers we are unable to examine the time course over which the impacts of the tsunami on thyroid hormone levels and health have emerged, beyond documenting their presence after 20 years. Future work with longitudinal biomarker measures can shed further light on the evolution of these outcomes. A second potential limitation may be the impacts of mortality selection at the time of the tsunami, if mortality due to the tsunami was related to worse health at the time. There are several reasons to believe that our results are not upwardly biased as a result of this concern. Mortality rates at the time of the tsunami were much higher among older populations – our observation of larger adverse tsunami impacts on health at older ages suggests that the results persist even among populations that faced the most mortality selection (Frankenberg et al. 2011). More generally, we would expect mortality selection on the basis of health to bias results toward zero – since we observe deleterious impacts of the tsunami. We find suggestive evidence for this in appendix figure 5. The non-parametric relationship between FT3 and baseline community mortality rates clearly shows the negative relationship that we report, but at the highest levels of baseline community mortality the relationship turns back slightly positive, potentially suggesting a scarring effect at lower mortality rates and a selection effect that starts to dominate at high ones. Nonetheless, our results are robust to these potential selection impacts at the highest levels.

In summary, these data provide rigorous new evidence for long-term changes in thyroid hormone physiology and cardiometabolic health after stressful events. Using a uniquely rich long-term panel study, we underscore the importance of long-term follow-up after stressful exposures: for many individuals, the long-term consequences of the 2004 Indian Ocean Tsunami on health are just beginning to emerge.

## Methods and data

### The Tsunami

The December 26, 2004 Sumatra-Andaman earthquake created an associated tsunami with waves up to 30 meters high, impacting communities along the western coast of Aceh, Indonesia’s western-most province. The tsunami killed roughly 5% of the population of Aceh, and displaced over half a million people.

At the time of the 2004 Indian Ocean tsunami no tsunami had affected mainland Sumatra for 600 years and no large-scale warning systems were in place. Substantial idiosyncratic local-area variation in tsunami impacts was caused by interactions between wave direction, seafloor, and land topography (Lavigne et al. 2009; McAdoo, Richardson, and Borrero 2007). It is, therefore, plausible to treat tsunami exposure within a *kecamatan* (sub-district) as exogenous to future health; evidence in support of this is presented in Table 1 and Figure 1. This exogeneity differs from many natural disasters – where early warnings or repeated occurrence results in selective avoidance behaviors so that exposure cannot be treated as exogenous (Baker 1991; Boustan, Kahn, and Rhode 2012; Currie and Rossin-Slater 2013).

### The Study of Tsunami Aftermath and Recovery

STAR is designed to provide causal evidence on the long-term impacts of the 2004 Indian Ocean tsunami (Frankenberg et al. 2008, 2011, 2025). Key strengths of the study design for this research are a natural-experiment centered on a large-scale sudden and unanticipated disaster that credibly identifies causal impacts, detailed and fine-grained information about exposures at the time of the tsunami, a population-representative sample, and successful longitudinal tracking over 20 years (achieving >93% follow-up including movers).

Other limitations in the literature that our study design addresses include reliance on opportunistic samples that are not population-representative and that in many cases are selected on health and well-being. As one example, movers after a disaster are seldom included in studies and yet movers are likely to be selected on health. Many studies rely on long-term recall about exposures that are potentially selective; and use measures that are not sufficiently fine-grained. (Aristizabal et al. 2020; Deryugina and Molitor 2020; Shiba et al. 2021).

The baseline, conducted in February and March of 2004, 10 months before the tsunami, was part of Statistics Indonesia’s annual national socioeconomic survey, SUSENAS, and is representative of the population living in coastal districts of Aceh, Indonesia. Baseline respondents have been tracked over time, including those who moved to other provinces. Follow-ups were conducted annually for the first 5 years post tsunami, and then 10, 13, 15, and 20 years post-tsunami.

The primary outcomes for this study are drawn from the 20-year follow up, fielded in 2024. The 20 year wave is focused on detailed health and cognitive assessments, following respondents who lived in a 25% random sub-sample of baseline communities. Thyroid measures were assessed for 3,270 individuals. Table 1 describes the sample.

UNC Campus IRB approved this project (protocol 20-2673).

### Biomarker outcomes

Cholesterols and FT3 were measured using gold-standard venous blood assays conducted in a validated clinical laboratory in Aceh, Indonesia. Hemoglobin A1c was assessed using an Afinion Point of Care Test in a climate-controlled setting, which we validated extensively. Body fat percentages were calculated using bioelectrical impedance analysis conducted with a Tanita scale. Blood pressures were assessed by trained field staff using manual sphygmomanometers.

We examine FT3, which we convert to a Z-scored outcome for comparability, and an inverse-covariance weighted cardio-metabolic health index calculated using the other biomarkers (Anderson 2008). This index also is standardized with mean 0 and standard deviation 1.

For a small subsample of individuals that overlap with a hair cortisol sub-sample collected 14 years after the tsunami, we also present descriptive results of the relationship between cortisol levels and free T3. Hair was collected on a random subsample of STAR respondents, and cortisol was ground and extracted as described in (Lawton et al. 2023). We use the natural logarithm of hair cortisol levels for analysis.

### Empirical approach

We draw comparisons between respondents living in communities in the same sub-district that experienced local differences in exposure to the 2004 Indian Ocean Tsunami, which we assess with a continuous measure of the mortality rate in a respondent’s community of residence (*desa*) *at the time of the tsunami*. These community-level differences are plausibly exogenous with respect to future health for two key reasons: 1) the tsunami was an unexpected, sudden shock. Sedimentary evidence suggests that mainland Sumatra had not seen a tsunami for over 600 years (Monecke et al. 2008). When surveyed at the pre-tsunami baseline, fewer than 10% of respondents reported their home faced any risk from a natural disaster, and fewer than 1% reported any risk of tsunami. These risks were also unrelated to eventual exposure. 2) the impacts of the tsunami on a given community depended on the interaction among topographical features of the land and shoreline, the seafloor, and the orientation of the coastline relative to the rift zone along which the earthquake occurred (Degueldre et al. 2016; Umitsu, Tanavud, and Patanakanog 2007). As shown in Figure 1, within a small geographic area, there was substantial variation in tsunami mortality. This variation is plausibly exogenous, and as shown in Table 1, not related to baseline characteristics or contemporaneous health behavior like smoking.

Exposure is assigned to respondents based on their community of residence at the time of the tsunami. To take into account that the distribution of mortality rates is right-skewed and include zeroes, we use a quartic root transformation in the models (which approximates a logarithmic transformation for positive numbers). We estimate the following linear regression for each health outcome, *Y*_*ick*_, conditional on sub-district (*kecamatan*) fixed-effects *γ*_*k*_, and covariates, *X*_*i*_. The covariates include education, height, a series age-by-sex interactions (implemented with 5 year age bins, to account for important age profiles in thyroid hormone levels by sex), current smoking behavior, and the elevation and distance to coastline of the community of residence at the time of the tsunami.

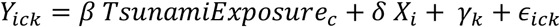

Standard errors are clustered at the level of the community of residence at the time of the tsunami.

We create a metabolic condition index that combines information from the cholesterol markers, %HbA1c, and blood pressures, which we use to aggregate outcomes and mitigate multiple testing concerns with several potential outcome variables. In order to create an index across multiple cardio-metabolic disease outcomes, we create an inverse-covariance weighted index of outcomes, with mean 0 and standard deviation 1 (Anderson 2008). We conduct heterogeneity analyses by age and sex with an extension of the above model, interacting *TsunamiExposure*_*c*_ with indicator variables for age group or sex.

In order to flexibly examine age-specific impacts of tsunami exposure in figure 2, we run a LOWESS-like procedure that estimates the impacts at each age. For each discrete age year, we apply triangular weights with a 10 year window on either side of the focal age group, and estimate the *β* coefficient that is specific to the age group from our primary specification. In table 2, we estimate heterogeneity by age groups. This is implemented with a single regression with interaction terms between the quartic root of fraction dead in the tsunami and a binary indicator for the demographic characteristic of interest.

## Acknowledgment

Financial support from the Broad Trauma Initiative (Subaward # 6910369-5500001914), the Burroughs Wellcome Fund Career Award at the Scientific Interface (Award #1018201), the National Institutes on Aging (R01AG031266, R01 AG065395, R24AG054365, and T32AG51108), the Eunice Kennedy Shriver National Institute for Child Health and Human Development (R01HD052762 and P2C HD050924), the National Institute of General Medical Sciences (T32GM144273), and the Wellcome Trust (OPOH 106853/A/15/Z) is gratefully acknowledged.

## Footnotes

1 Some cited studies use total T3 and some use free T3 – when describing other work, we summarize both as changes in T3. While closely related, free T3 is the most direct measure of active circulating T3 hormone. Prior work suggests total T3 follows similar age, demographic, and health patterns as free T3, but with attenuated signal (Lawton, Sabatini, and Hochbaum 2024).

2 The NHANES does not include specific month-of-interview, but does include a May-October indicator, which we use to define measurement time of year.

3 The monthly average high temperature in Aceh fluctuates between 86 and 90F during the year; in the US, it fluctuates between 67 and 85F between May and October. In NHANES, average FT3 levels are 0.1 standard deviations higher in the other months; free T4 levels are 0.03 standard deviations higher (not statistically significant) and TSH levels are the same, conditional on age, sex, and race (Lawton et al. 2024).

4 For binary classifications, we use a damage zone classification that stratifies communities into those heavily damaged versus not heavily damaged, based on information sources that include satellite imagery, community informant reports collected after the tsunami, and the observations of interviewers working in the field after the tsunami. (Frankenberg et al. 2008)

5 In the figures, the NHANES sample is restricted to those measured in May-Oct and in the age figure, the NHANES sample is restricted to non-obese (BMI<30) individuals to mitigate body composition differences across samples. These adjustments substantially affect the concordance between the NHANES and STAR demographic patterns. The NHANES FT3-age slopes are roughly 30% steeper, and FT3-BMI slopes for males are 50% steeper, without these adjustments.

6 Mortality is considered tsunami-related if it occurred within 5 days after the tsunami.

7 Demographic comparisons calculated in the STAR population

8 We also explored a measure of exposure that stratifies communities that were heavily damaged, moderately damaged and not damaged, based on information from high-resolution satellite imagery, community informants, and STAR interviewer observations (Frankenberg et al. 2011). The estimated impact of living in a heavily-damaged community is large (−0.1 standard deviations) but only statistically significant at a 9% size of test in models with district fixed effects (Appendix Table 2). Estimates are the same but not significant with sub-district fixed effects because there is little variation in damage within sub-district. This highlights the advantage of using continuous measures of exposure and identifying dose-response effects. The stratification is useful to identify those who were not directly impacted by the tsunami, in order to draw comparisons with the NHANES.

