## Supplement for "Stressful exposure from the 2004 Indian Ocean Tsunami drives long-term changes in thyroid hormone physiology"

**Appendix**

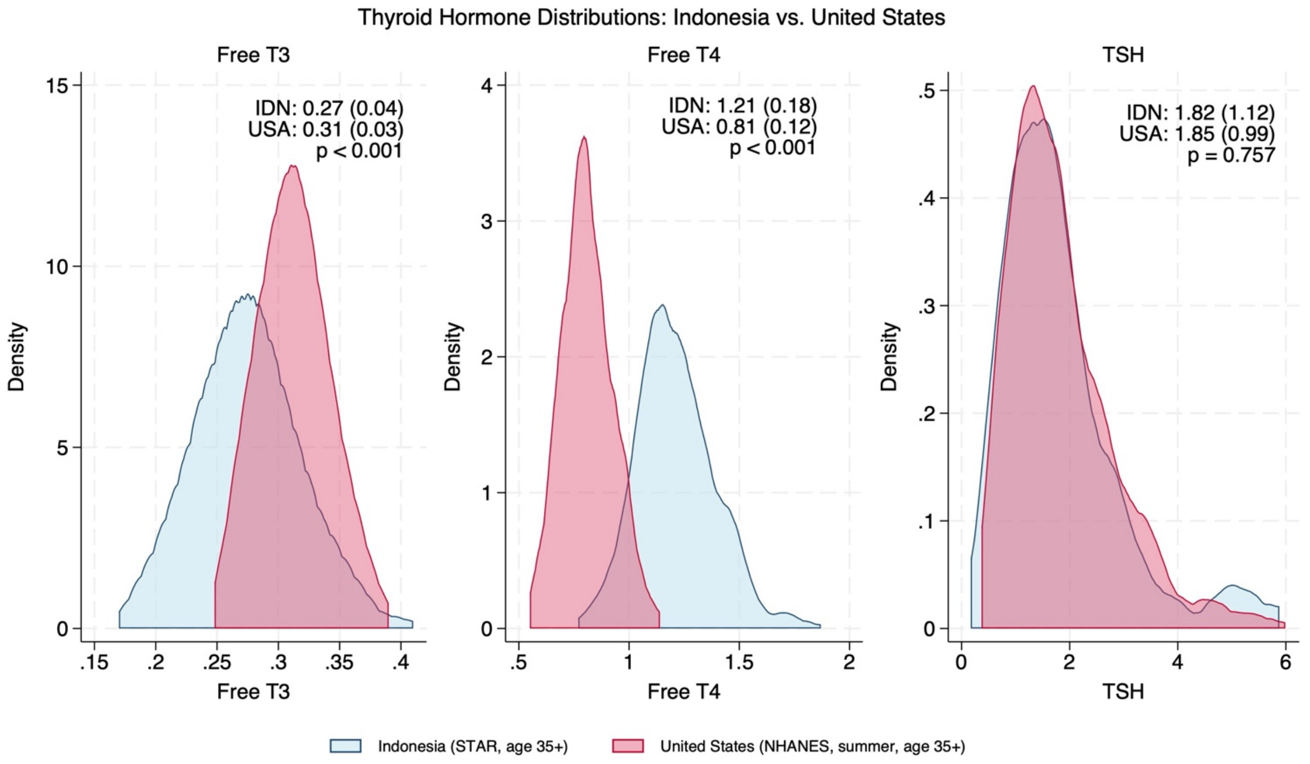


**Appendix Figure 1.** Kernel density plots of thyroid hormone distributions in Indonesia (data from STAR) and from the US (data from the NHANES 2011-2012 wave). US data have been age-adjusted to match the age distribution of STAR. Means and standard deviations reported on the figure, as well as p-values for the difference in means between the NHANES and STAR. STAR collected FT3 measures on 3,270 individuals, free T4 on 378, and TSH on 183 individuals.

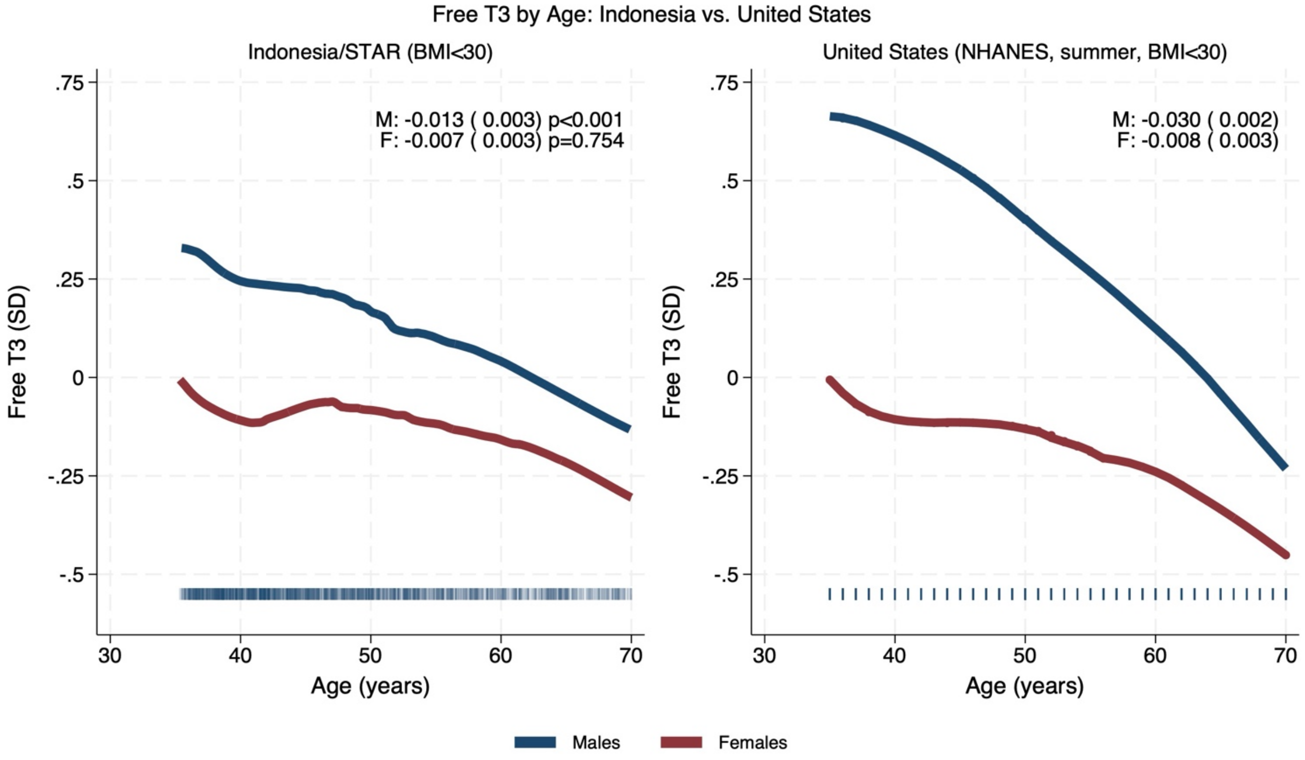


**Appendix figure 2:** Non-parametric relationships between FT3 and age in the STAR and NHANES studies. Linear estimates of the sex-specific age slopes are included, as are the p-values for a test of the null that slopes are the same in the US and in Indonesia. To minimize confounding by tsunami exposure, we conservatively limit the STAR sample to communities with less than 5% fraction dead that were also not heavily exposed to tsunami damage. Markers along the bottom show the distribution of observations along the x-axis variable. For the NHANES, only age in years is observed.

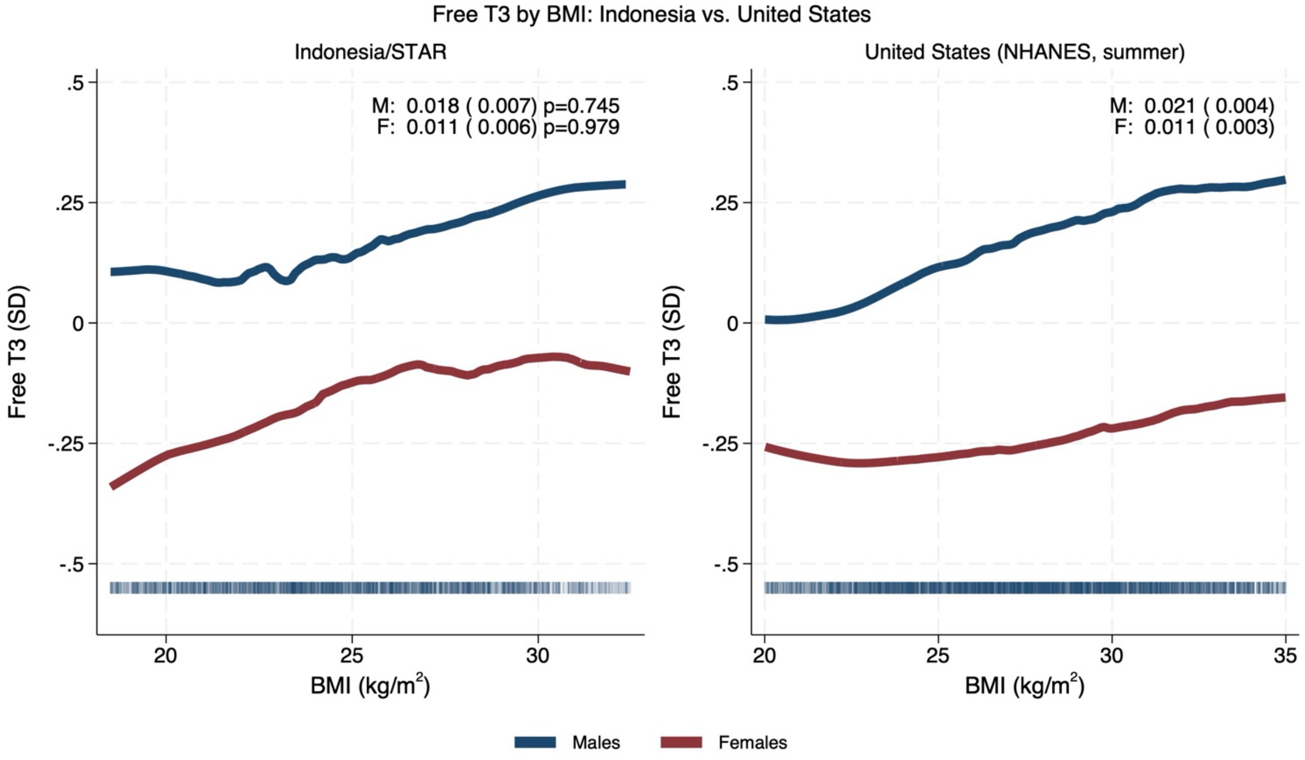


**Appendix figure 3:** Non-parametric relationships between FT3 and BMI in the STAR and NHANES studies. Linear estimates of the sex-specific BMI slopes are included, as are the p-values for a test of the null that slopes are the same in the US and in Indonesia. To minimize confounding by tsunami exposure, we conservatively limit the STAR sample to communities with less than 5% fraction dead that were also not heavily exposed to tsunami damage. In this figure, we use BMI rather than our preferred measure of fat percent, so that we can draw harmonized comparisons with the US. Markers along the bottom show the distribution of observations along the x-axis variable.

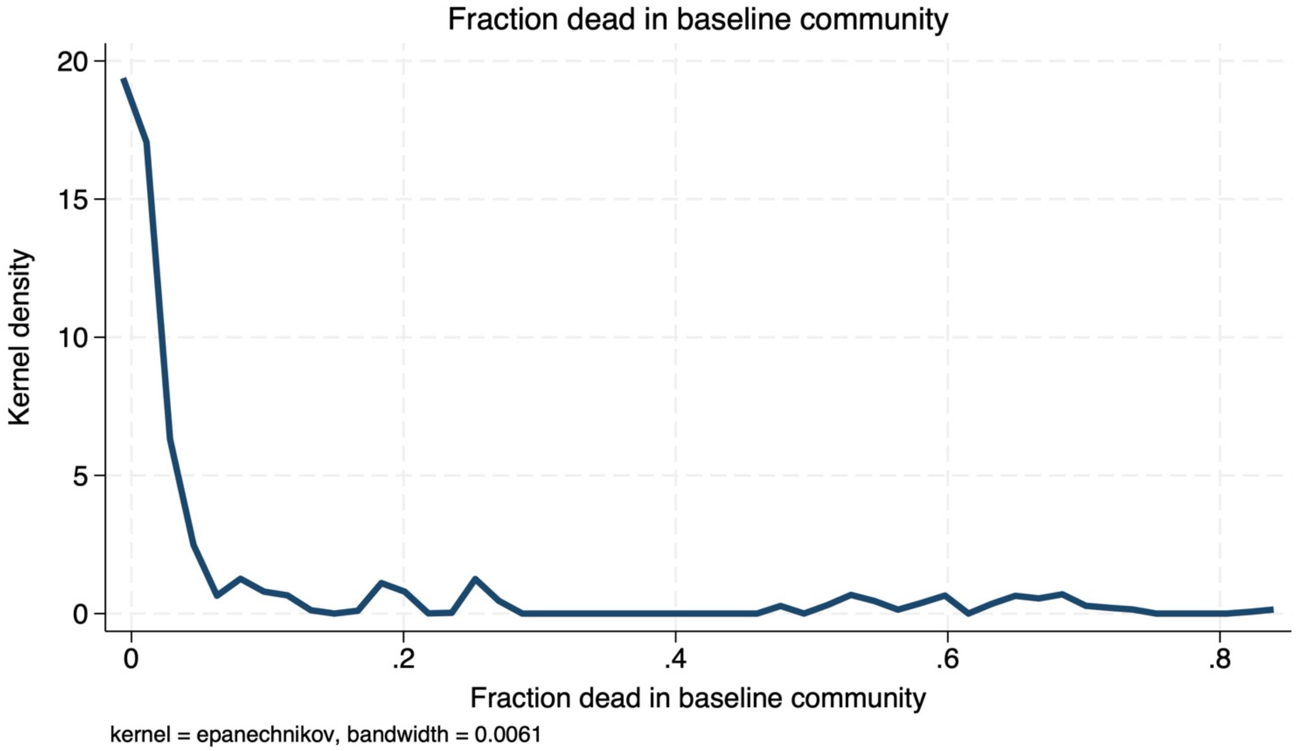


**Appendix figure 4:** Kernel density of mortality rates in STAR communities at the time of the tsunami.

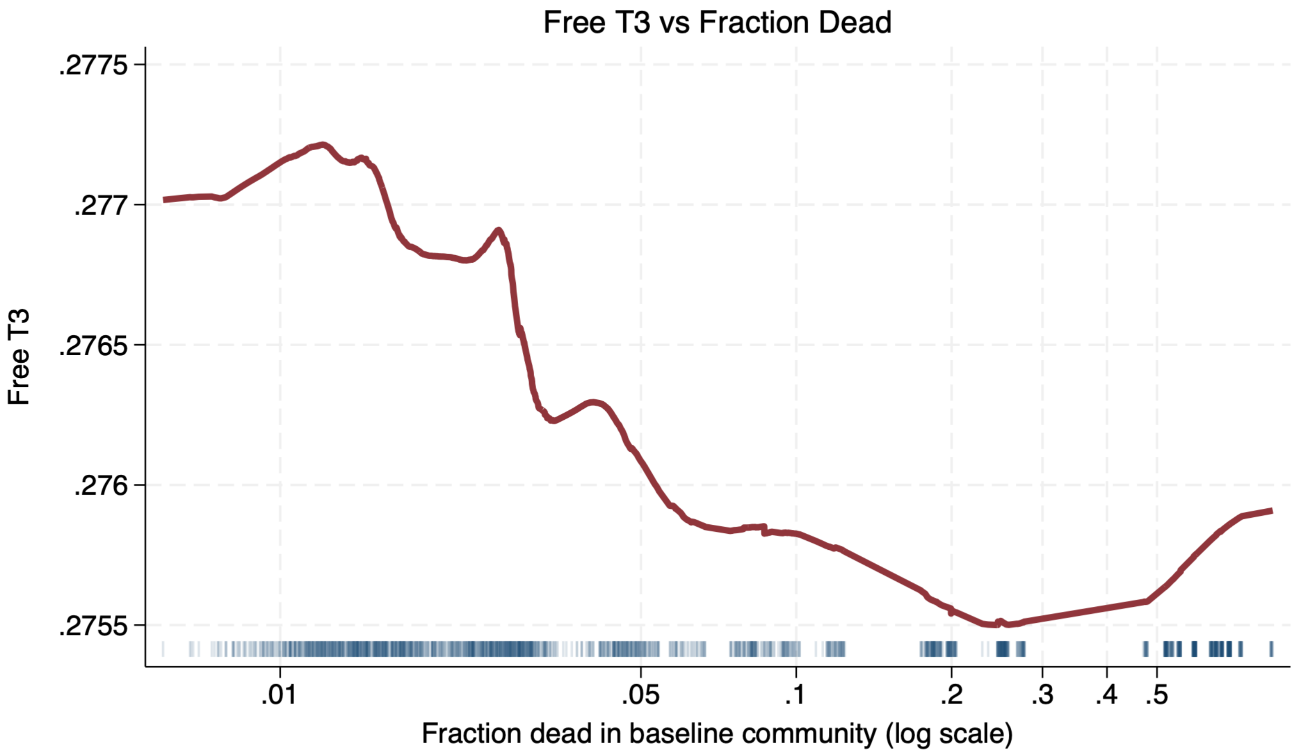


**Appendix figure 5:** LOWESS estimate of FT3 vs fraction dead in the tsunami in the baseline community. Markers along the bottom show the distribution of observations along the x-axis variable.

### Appendix Table 1. Coefficients for metabolic index sub-components

| | [1] | [2] |
|---|---|---|
| | Coefficient on fraction dead | Coefficient on fraction roads damaged |
| Variable | Coefficient (SE) | Coefficient (SE) |
| Free T3 | -0.278 | -0.138 |
| | ( 0.137) | ( 0.058) |
| Metabolic Index | 0.308 | 0.097 |
| | ( 0.082) | ( 0.048) |
| HDL Cholesterol | -0.232 | -0.016 |
| | ( 0.119) | ( 0.063) |
| Non-HDL Cholesterol | 0.34 | 0.166 |
| | ( 0.127) | ( 0.121) |
| HbA1c | 0.068 | 0.06 |
| | ( 0.089) | ( 0.045) |
| Systolic BP | 0.103 | -0.101 |
| | ( 0.093) | ( 0.054) |
| Diastolic BP | 0.152 | 0.055 |
| | ( 0.137) | ( 0.042) |

Notes: STAR analytical sample, 3,270 observations. Point estimates are presented with clustered standard errors from a regression of the quartic root of fraction dead in the tsunami, and a regression on fraction of roads damaged by the tsunami. Regressions conditional on kecamatan (sub-district) fixed effects, elevation, distance to the coastline, 5 year age bin-by-sex interactions, smoking, and years of education. Standard errors are clustered at the baseline community-of-residence.

### Appendix Table 2. Coefficients on binary damage zone classifications on free T3

| | [1] | [2] |
|---|---|---|
| | Coefficient on heavy damage zone | |
| | With district FE | With sub-district FE |
| Variable | Coefficient (SE) | Coefficient (SE) |
| Free T3 | -0.102 | -0.115 |
| | ( 0.060) | ( 0.125) |

Notes: STAR analytical sample, 3,270 observations. Point estimates are presented with clustered standard errors from a regression of the quartic root of fraction dead in the tsunami. Regressions conditional on specified fixed-effects, 5 year age bin-by-sex interactions, smoking, and years of education. Standard errors are clustered at the baseline community-of-residence.
